# Prognostics for pain in osteoarthritis: Can clinical measures predict pain after total joint replacement?

**DOI:** 10.1101/751503

**Authors:** Joana Barroso, Kenta Wakaizumi, Diane Reckziegel, João Pinto-Ramos, Thomas Schnitzer, Vasco Galhardo, A. Vania Apkarian

## Abstract

A significant proportion of osteoarthritis (OA) patients continues to experience moderate to severe pain after total joint replacement (TJR). So far, preoperative factors related to pain persistence have been mainly studied using individual predictor variables and distinct pain outcomes, thus leading to lack of consensus in the field. In this prospective observational study, we evaluated knee and hip OA patients before, 3 and 6 months post-TJR searching for clinical predictors of pain persistence. We assessed multiple measures of quality, mood, affect, health and quality of life, together with radiographic evaluation and performance-based tasks, modeling four distinct pain outcomes. Multivariate regression models were built, and a network analysis was applied to pain related biopsychosocial measures and their change with surgery. A total of 106 patients completed the study. Pre-surgical pain levels were not related to post-surgical residual pain. Distinct pain scales were associated with different aspects of the pain experience. Multi-factorial models did not reliably predict post-surgical pain in knee OA across four distinct pain scales and did not generalize to hip OA; however, network analysis of pain related biopsychosocial measures showed significant changes post-surgery in both groups. Our results show that although tested clinical and biopsychosocial variables are reorganizing after TJR in OA, they do not present as a robust markers for post-surgery pain outcomes. A better understanding of mechanisms underlying pain persistence after TJR is necessary to derive clinical prognostic factors.

## Introduction

Osteoarthritis is the most common cause of arthritis worldwide and a major cause of chronic musculoskeletal pain. Although nociceptive inputs elicited by joint degeneration and chronic inflammation are commonly recognized as the main contributing factors, current understanding of OA pain pathophysiology remains incomplete. In the last few years, a growing body of research indicates that altered peripheral and central nociceptive processes are influential [1]. This is substantiated by the discordance in joint structural damage and pain intensity [2], but also by the results of surgical treatment [3]. Total joint replacement (TJR) is regarded as an effective and safe intervention for advanced hip and knee OA; nevertheless, an important proportion of patients still report moderate to severe persistent pain post-TJR, not attributable to identifiable surgical or clinical complications. In the case of knee OA (KOA), persistent post-surgical pain is reported in about 20% of the patients. For hip OA (HOA), this number appears to be lower (up to 10%) [4].

Although considerably studied, persistent post-TJR pain is not completely understood. Contrary to other types of chronic post-surgical pain, defined as pain that occurs or intensifies after a surgical procedure and lasts for at least 3 months [5], in OA, pain is a pre-existing condition, leading to a less clear delineation of unsuccessful outcomes. This reflects in distinct definitions of persistent pain across studies [6].

Regarding risk factors for pain persistence after TJR, those have been proposed, mainly for KOA [7]. Pain intensity prior to surgery, disproportion between pain intensity and articular damage, neuropathic-like symptoms, psychosocial factors such as pain catastrophizing and poor coping strategies [8] are commonly referenced as important predictive factors. Although these have been studied repeatedly, an extensive variation of outcome measures can be found and there is no agreement on which measures should be used to assess chronic pain after TJR [7]. The proposed risk factors across studies are often diverse, tested thru univariable associations, different study designs and analysis methods. Overall, the quality of evidence on prognostic factors for recovery after total knee replacement (TKA) is low [9].

Here, in a prospective cohort study, we purpose to study how distinct pain measurement instruments relate to different aspects of pain in OA. We aim to develop and evaluate models predictive of pain and pain relief after surgery for knee and hip OA pain. Additionally, by using a global network analysis approach, we purpose to assess the reorganization of pain related clinical and biopsychosocial properties of KOA and HOA patients after TJR.

## 2. Materials and Methods

### 2.1 Study sample

KOA and HOA patients with clinical indications for primary arthroplasty surgery participated in this longitudinal observational study. The present report is part of a brain neuroimaging study, studying central mechanisms in osteoarthritis, which will be latter reported.

Enrollment took place at the Orthopedic Surgery Department of Centro *Hospitalar de São João*, a tertiary care hospital in Porto, Portugal. Study protocol was approved by the local Ethics Committee, and all participants provided informed written consent prior to partaking in the study. Sample size was determined by the number of patients waiting for surgery who met the eligibility criteria for the study, during a period of 20 months. Initial evaluation happened 1-3 months before TJR surgery and follow-up continued up to 6 months after surgery. A total of 95 knee OA and 25 hip OA patients, and 37 healthy control subjects were included (the latest group not studied in this report).

Eligible patients met the following inclusion criteria: age between 45 and 75 years-old; diagnosis of HOA and KOA according to the clinical classification criteria of the *American College of Rheumatology*, and surgical indications for TJR (criteria for surgery selection was moderate to severe pain and quality of life impairment, after clinical and radiological evaluation and medical decision by a certified orthopedic surgeon in our center). Patients were excluded when there was evidence of secondary OA due to congenital or development diseases and inflammatory bone and articular diseases. Bilateral OA with predicted indication for contralateral arthroplasty in the following year, other chronic pain conditions (e.g., fibromyalgia; chronic pelvic pain) and chronic neurological or psychiatric disease (e.g., depression major, dementia, obsessive compulsive disorders, Parkinson’s disease, demyelinating diseases, peripheral sensory neuropathy), were also exclusion criteria, as well as cognitive impairment. Previous history of stroke or traumatic brain injury was also exclusionary. Secondary OA following history of minor trauma or previous arthroscopy surgery due to ligamentous/meniscal injury was not an exclusion criterion.

### 2.2 Study design

This study comprised a total of 4 visits. Patients were initially assessed 1-3 months before surgery (V1). A second pre-surgical visit was held 2 to 6 weeks prior surgery at the neuroimaging facility (V2). Two post-surgical visits (V3-V4) occurred at 3 months and 6 months post-surgery. Specific data collected at each visit are shown in figure 1A. During visits 1, 3 and 4 patients were assessed for: (1) Clinical and socio-demographic properties; (2) physical function – performance-based tests; (3) radiographic evaluation, (4) pain, mood and health questionnaires; brain imaging was performed at visits 2 and 4.

**Figure 1A.**
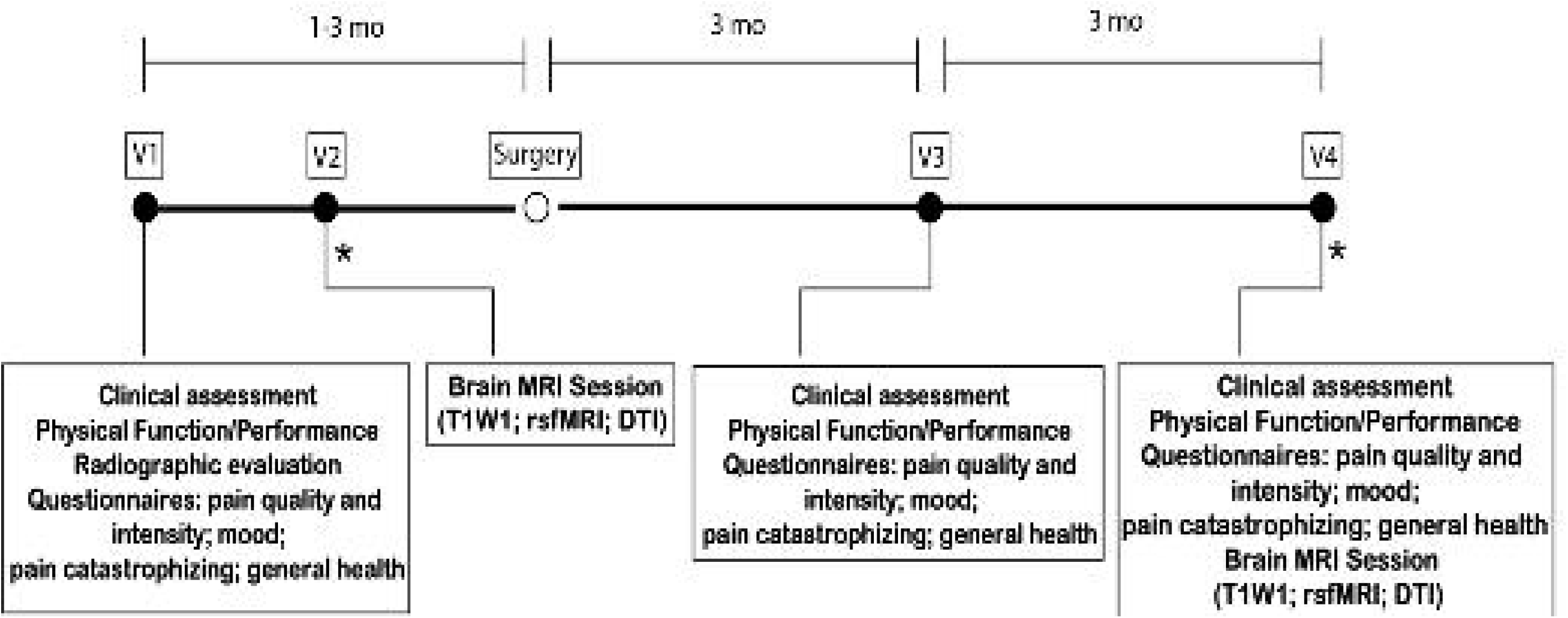
Experimental design, timeline and data collected. Knee and hip osteoarthritis patients entered a 4 visit (V1-V4), pre- and post-total joint replacement surgery, longitudinal, observational study. V1 and V2 occurred before surgery. V3 and V4 took place at 3 and 6 months after surgery. At each visit, participants underwent a series of assessments. *Brain MRI session. mo, months; MRI, magnetic resonance imaging; T1WI, T1-weighted imaging; rsfMRI, resting state functional magnetic resonance imaging; DTI, diffusion tensor imaging.

### 2.3 Measures

#### 2.3.1 Clinical and demographic assessment

Demographic profiling, acquired at V1, included age, education and professional status. Medical data concerning height and weight, pre-surgical co-morbid conditions, previous surgeries, general medication and smoking habits were recorded at patient interview and by clinical charts analysis. A clinical questionnaire regarding the history and evolution of knee pain assessed pain onset, duration and frequency; pain medication and previous non-pharmacological treatments. The Medicine Quantification Scale (MQS) was used to score type and dose of pain medication [10]

At the post-surgical visits (V3-V4) a second clinical questionnaire assessing pain recovery, time to recovery, patient satisfaction, pain medication, use of health care services and rehabilitation protocol was administered.

#### 2.3.2 Physical function – performance-based tests

Physical function was assessed with two different tests, depending on the activity measured. Ambulatory transitions were evaluated with the Timed up and go test (TUG) [11], and aerobic capacity/walking long distances with the six-minute walk test (6MWT) [12-14]. These tests were selected based on the OARSI 2013 recommendations [15].

#### 2.3.3 Radiographic assessment

As part of standard hospital protocol, patients scheduled for TJR had bilateral joint radiographs during the 6 months before surgery. Knee OA radiographs were taken in two views: anterior-posterior (AP) weight-bearing with knee flexion at 20° and foot internal rotation at 5°, and horizontal beam lateral view, with lateromedial projection, the patient in supine position and the knee flexed at 30°. Hip OA patients had AP supine radiograph of the pelvis, with lower limbs internally rotated 15° degrees from the hip.

Radiographs were scored accordingly to the Kellgreen-Lawrence (KL) classification - grades 0 to 4 [16], by two trained radiologists. The first classified the whole sample, the second classified half of the subjects for inter-reliability measurement. Both researchers were blind to the clinical data of the patients when scoring. Inter-rater reliability was determined for KOA imaging only and the intra-class correlation coefficient of KL grading was 0.91 (95% confidence interval 0.80-0.93).

#### 2.3.4 Questionnaires – Pain, Mood and Health

Seven questionnaires were administered by a trained clinician, during face-to-face interview. They were administered both before surgery (V1), and in the post-surgical visits (V3-V4). The repeated use of the same measures allowed us to track changes concerning intensity and quality of pain, emotion and affect, health and quality of life. All questionnaires were used in their validated Portuguese version. We assessed: 1) KOOS, HOOS, validated injury and OA outcome scores for knee and hip [17-19]; 2) Brief Pain Inventory – Short Form (BPI) [20-22]; 3) McGill Pain Questionnaire (MPQ) [23, 24]; 4) Doleur Neuropathique en 4 Questions (DN4) [22, 25]; 5) Hospital Anxiety and Depression Scale (HADS) [26, 27]; 6) Pain Catastrophizing Scale (PCS) [22, 28]; and 7) SF36-item Short Form Survey (SF36) [29, 30].

### 2.4 Primary outcome variables

Primary outcome variables were part of the questionnaires/clinical assessments and consisted of 4 distinct pain intensity related scales/subscales: Numeric Rate Scale (NRS); BPI – Pain Severity; KOOS Pain and HOOS Pain, as clinically appropriate; SF36 Bodily Pain, here addressed specifically for knee/hip articular pain.

For each of the 4 outcome measures and for an aggregate of all four, we examined relationships for pain relief post-surgery on a per subject basis, by calculating residual pain: %residual pain = 100 – (100 *(average pain pre-surgery - post-surgery pain [at 3, or 6, months])/ average pain pre-surgery)). Thus, 100% residual pain = no change in a given pain measure between before and after surgery; 0% residual pain = complete relief from initial pain; while values >100% indicate worsening of pain post-surgery.

### 2.5 Statistical analysis

All data from the reported measures were manually entered by the same researcher. Regarding missing data, when at least 30% was missing from a questionnaire (total or sub-score if applicable), it was excluded. When missing data were less than the threshold 30%, we used the mean of the total score/sub-score to fill in missing items.

Descriptive statistics were used to describe the study sample, with continuous variables presented as mean and standard deviations and categorical data as numbers and percentages. Comparisons between the two OA groups used independent sample t-tests or Chi-square(*X*^*2*^) tests, for continuous parametrical variables and categorical data respectively.

Interrelationship of the primary outcome variables (all scored on a 0-10 score) was assessed through correlation analysis using Pearson product-moment tests. The effects of time (pre-, 3- and 6-months post-surgery), type of OA and pain outcome measure on pain intensity were studied with a three-way mixed ANOVA. Following the initial procedure, two-way interactions and simple main effects were considered and pairwise comparisons with Bonferroni adjustments were performed.

A data dimensionality reduction from all 19 subscales of 7 questionnaires and 2 physical performance scores was achieved using a principal component analysis (PCA) in KOA patients at baseline. Overall and individual Kaiser-Meyer-Olkin measures were 0.86 and >0.5 respectively. Threshold for component retention was set on eigenvalues >1.0, together with visual inspection of the scree plot for evaluation of the inflection point. A factor rotation on the obtained components was applied using a Promax oblique rotation technique. Threshold of factor loading was set on 0.5/-0.5 and components were labeled given the observed loadings. Due to the limited number of subjects available in the HOA group, we generated the component values using the same weights retrieved with PCA for the KOA, which enables direct comparison of TJR effects on network properties.

Different regression analysis techniques were used to model pain outcomes in KOA and HOA. For KOA, multifactorial regression models were generated using a stepwise forward and backward selection method, in an automatic step-by-step iterative construction of the model.

Significance level to enter (α-to-enter) was set at 0.05 and α-to-remove at 0.10. To test if the models obtained in KOA replicated in HOA patients, and due to a smaller sample size in this group, we applied a multiple linear regression analysis in HOA, entering as independent variables the predictor factors uncovered for KOA, thus testing the extent of shared factors between the two conditions. For all regression models, assumptions of linearity, independence of observations, homoscedasticity and absence of multicollinearity were met, and residuals were approximately normally distributed in all models. Outliers were detected by examining studentized deleted residuals, any values greater than ± 3 standard deviations were removed. Throughout all models, no more than 3 cases were removed.

A composite measure of pain intensity was built averaging the four outcome scales. Here, a two-way mixed effects ANOVA was conducted to study differences in pain levels across time and type of OA. The same regression analysis methodology was applied to this new variable in HOA and KOA groups.

Correlation matrices of the clinical and psychological variables (questionnaires subscales and physical performance scores) were represented as binarized networks, constructed at the 25% stronger correlations for each matrix (KOA/HOA at baseline, 3- and 6-months post-surgery), and visualized using the software Cystoscape (v3.6.1, http://www.cytoscape.org). For each network, questionnaire measures were represented as nodes and the thresholded correlations as edges. Network communities were derived from the previous PCA. Two network graph measures were computed to characterize and quantify topological changes, using the Matlab Brain Connectivity Toolbox [31]. Clustering coefficient is a measure of the extent to which nodes in a graph tend to cluster together. Nodes have the trend to create groups characterized by a high density of connections. We computed local clustering coefficient of all nodes, and averaged them, reflecting the overall level of clustering in a network, from 0 (no clustering) to 1 (maximal clustering). The second calculated measure, modularity, refers to the compartmentalization and interrelation of modules in a network. Modules can be defined as sets of nodes densely connected among themselves and poorly connected to other regions of the network. Using the Louvain community detection algorithm, averaged over 100 computed repetitions, we obtained values that vary from 0 (random network) and 1 (highly structured network).

We studied the changes in the strength of connectivity for all networks, calculating the change in correlation coefficients for all pairs of subscales from baseline to three and six months, and averaging these over the entire networks, obtaining the mean ΔR. For all inter and intra-group comparisons, regarding network measures and change in correlation coefficients, statistical probability was computed with 10,000 repeated random resampling.

All data were analyzed using the Statistical Package for the Social Sciences (IBM Corp. Released 2016. IBM SPSS Statistics for Windows, Version 24.0. Armonk, NY: IBM Corp), JMP software (JMP^®^, Version *14*. SAS Institute Inc., Cary, NC, 1989-2007) and MATLAB (MATLAB and Brain Connectivity Toolbox release 2016a, The Mathworks, Inc., Natick, Massachusetts, US).

## Results

### 3.1 Recruitment, assessment and participant characteristics

A total of 94 KOA and 25 HOA patients were eligible and agreed to participate in this longitudinal, observational study. At 6 months, a total of 84 KOA and 22 HOA completed the study. **Figure 1B** presents patient and control participants flowchart and timeline. Causes for withdrawal included: revision arthroplasty due to periprosthetic infection or prosthesis displacement (n=4); other co-morbidities (n=2, concomitant oncological disease) and voluntary withdrawal (n=12).

**Figure 1B.**
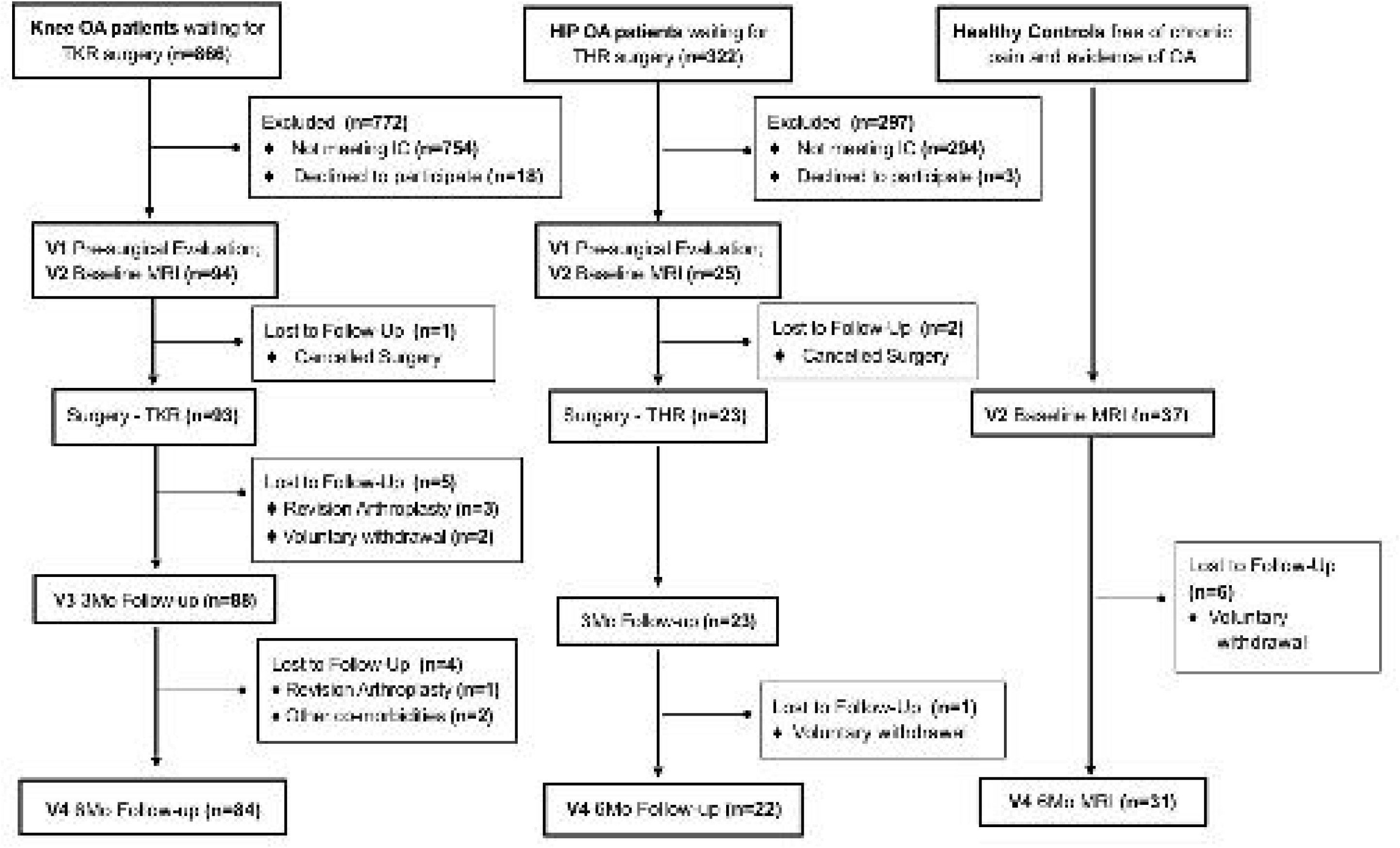
Recruitment and retention for KOA and HOA, and healthy control participants. The full battery of assessments was performed in osteoarthritis patients. Healthy individuals were recruited to act as controls in brain imaging analyses (not reported here). All patients were recruited from the same tertiary care hospital. Healthy participants were recruited from the general population in the Porto area.

**Table 1** describes HOA and KOA patients’ demographic characteristics. Mean age of KOA patients was greater than that of HOA patients; the KOA group was predominantly female while the HOA group included mostly males. Body mass index (BMI) was higher in KOA than HOA patients. Smoking habits, educational level and habitation status were similar between HOA and KOA. Regarding occupational status, for the KOA group the most common status was retirement; HOA patients were mainly on medical leave, which relates to their differences in age.

**Table 1.**
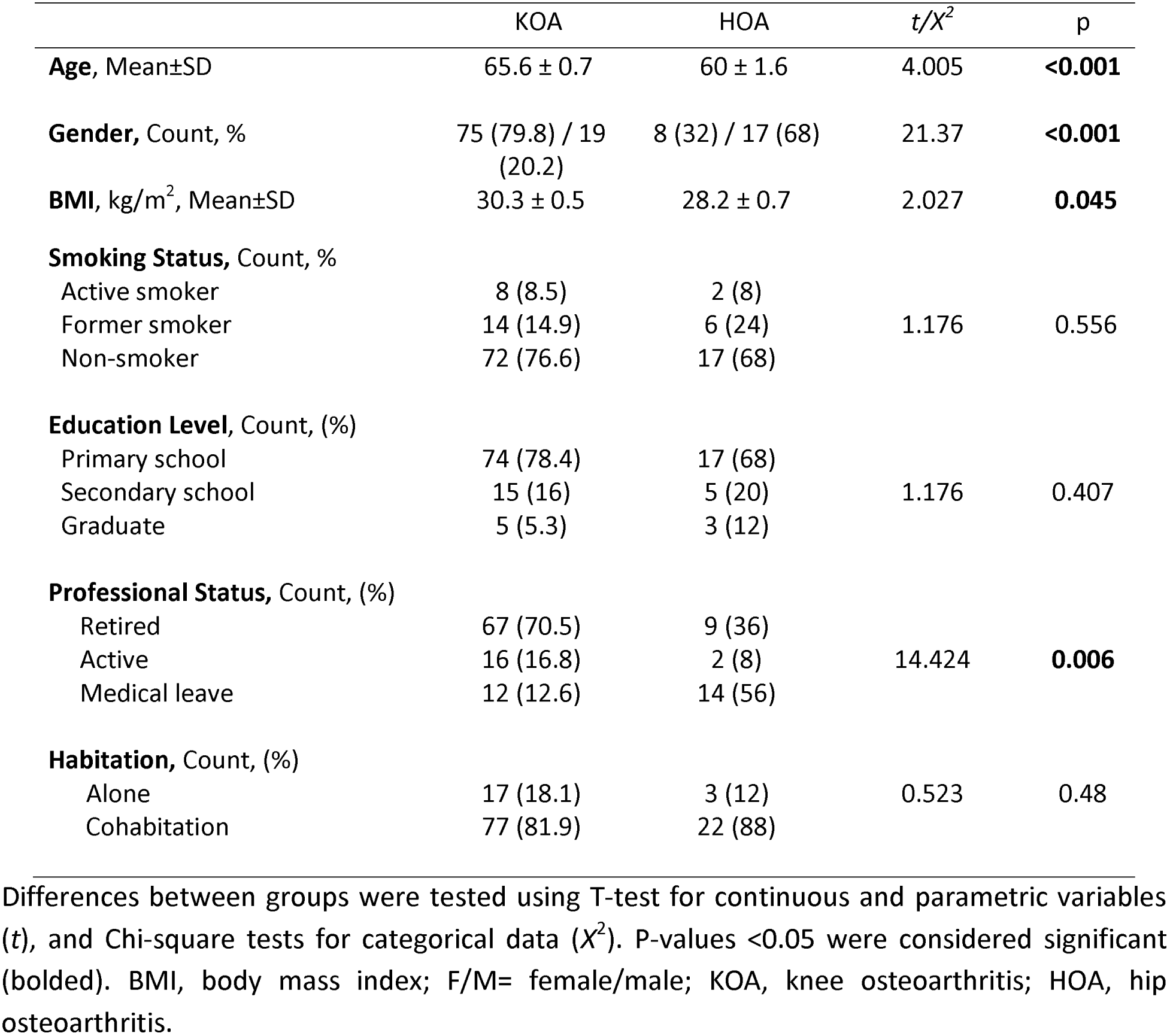
Demographic characteristics of KOA and HOA patients.

IC, Inclusion criteria; MRI, magnetic resonance imaging; KOA, knee osteoarthritis; HOA, hip osteoarthritis; THR, total hip replacement; TKR, total knee replacement.

### 3.2 Pain intensity as a function of type of pain measurement instrument, surgery, time, and OA joint involvement

We examined the pain intensity determined by our four pain outcome measures (NRS, KOOS pain, BPI pain severity, SF-36 pain), both at baseline and after surgery, in KOA and HOA patients, and then evaluated their interrelationship **(Table 2)**. All pain magnitudes decreased post-surgery, correlations among measures generally strengthened. Mean post-surgical pain levels (across all measures) was lower in the HOA group than in the KOA group, and the pain intensity estimate was highest with the SF-36 pain scale.

**Table 2.** Pain in KOA and HOA patients pre- and post-surgery, characterized with four outcome measures.

| Pain Intensity-Related Measure |  |  |  |  |  |
| --- | --- | --- | --- | --- | --- |
| Baseline |  |  |  |  |  |
| Knee OA (n=84) | M | SD | (1) | (2) | (3) |
| (1) BPI Pain Severity | 4.79 | 1.5 |  |  |  |
| (2) NRS | 6.53 | 1.67 | .792** |  |  |
| (3) KOOS Pain | 6.49 | 1.49 | .266** | .313** |  |
| (4) SF-36 Pain | 7.04 | 1.82 | .162* | .288* | .475** |
| Hip OA (n=22) |  |  |  |  |  |
| (1) BPI Pain Severity | 4.38 | 1.52 |  |  |  |
| (2) NRS | 6.09 | 1.66 | .801** |  |  |
| (3) HOOS Pain | 5.86 | 1.63 | .564* | .405 |  |
| (4) SF-36 Pain | 6.49 | 1.55 | .547* | .474* | .631* |
| 3 Months |  |  |  |  |  |
| Knee OA (n=84) | M | SD | (1) | (2) | (3) |
| (1) BPI Pain Severity | 1.69 | 1.48 |  |  |  |
| (2) NRS | 1.89 | 2.03 | .922**<sup>†</sup> |  |  |
| (3) KOOS Pain | 2.45 | 1.96 | .837**<sup>†</sup> | .809**<sup>†</sup> |  |
| (4) SF-36 Pain | 3.39 | 2.25 | .750**<sup>†</sup> | .771**<sup>†</sup> | .774**<sup>†</sup> |
| Hip OA (n=22) |  |  |  |  |  |
| (1) BPI Pain Severity | 0.54 | 1.05 |  |  |  |
| (2) NRS | 0.55 | 1.06 | .920** |  |  |
| (3) HOOS Pain | 0.79 | 0.88 | .257 | .159 |  |
| (4) SF-36 Pain | 1.95 | 1.71 | .329 | .356* | .381* |
| 6 Months |  |  |  |  |  |
| Knee OA (n=84) | M | SD | (1) | (2) | (3) |
| (1) BPI Pain Severity | 1.70 | 1.53 |  |  |  |
| (2) NRS | 2.01 | 1.9 | .930**<sup>†</sup> |  |  |
| (3) KOOS Pain | 2.19 | 2.17 | .813**<sup>†</sup> | .784**<sup>†</sup> |  |
| (4) SF-36 Pain | 2.98 | 2.26 | .671*<sup>†</sup> | .652**<sup>†</sup> | .791**<sup>†</sup> |
| Hip OA (n=22) |  |  |  |  |  |
| (1) BPI Pain Severity | 0.63 | 0.76 |  |  |  |
| (2) NRS | 0.73 | 0.93 | .961**<sup>†</sup> |  |  |
| (3) HOOS Pain | 0.80 | 0.77 | .379 | .280* |  |
| (4) SF-36 Pain | 1.98 | 1.87 | .471* | .479* | .559** |

Four scales were used (BPI Pain Severity, NRS, HOOS Pain, SF-36 Pain; all presented on a 0-10 scale) to assess pain pre-surgery (Baseline) and 3, and 6 months post-surgery. Mean (M), standard deviation (SD) and Pearson's product-moment correlations between the 4 scales are presented. For both OA groups the four measures decrease in amplitude after surgery and are correlated with each other, improving following surgery in the KOA group. \* $p < 0.05$ ; \*\* $p < 0.01$ . <sup>†</sup>Significant increase in correlation from baseline, $p < 0.05$ . KOA, knee osteoarthritis; HOA, hip osteoarthritis; BPI Severity, Brief Pain Inventory Pain: severity subscale; HOOS Pain, Hip Injury and Osteoarthritis Outcome Score: pain subscale; NRS, Numeric Rating Scale; SF36 Pain, Short-form (36) Health Survey: pain subscale.

A three-way ANOVA was conducted to determine the effects of time (pre-, 3, 6, months post-surgery), the four pain outcome measures, and the type of joint OA, on pain intensity. We found a non-significant three-way interaction between these variables. The two-way interactions were statistically significant between pain measures and time (F(6,624)=5.231, p<0.001); type of OA and time (F(2,624)=4.096, p=0.022); but not type of OA and types of pain measures (F(3,624)=2.021, p=0.129) revealing that questionnaires show a similar rating pattern for hip and knee OA.

Main effects of pain measurement types were statistically significant at baseline, 3 and 6 months (F (3,315) =66.6, p<0.001; F (3,315) =51.03, p<0.001; F (3,315) =41.02, p<0.001). Pairwise comparisons revealed that at baseline, pain intensity estimates were lowest for BPI pain severity, (mean differences - NRS: −1.72 [-2.44, -0.99], KOOS/HOOS: −1.436 [-2.158, -0.71], SF36: −2.06 [−2.78, −1.4], p<0.001). At 3 and 6 months after surgery pain intensity was higher when measured by SF-36 pain (mean differences at 3 months: NRS: 1.269 [0.509,2.03], BPI:1.566 [0.8,2.33], KOOS [0.24,1.76], p<0.003; at 6 months: NRS: 1.038 [0.23,1,81], BPI: 1.354 [0.57,2.14], p<0.003).

Joint involvement was also significant: KOA patients had higher levels of reported pain at baseline that HOA patients (mean difference: 0.55 [0.17,0.93], p=0.005), while HOA surgery resulted in a larger decrease in pain intensity than KOA surgery (mean differences at 3 months: 1.462 [1.06,1.87] and 6 months: 1.21 [0.8,1.62], p value <0.001).

The main effect of time on pain intensity showed that from baseline to 3 months there is a large decrease in pain intensity. There was no change in pain intensity between 3 months and 6 months, revealing that pain levels were stable from 3 months onwards in both OA groups (mean differences, KOA: Baseline-3 months 3.8 [3.512,4.124] p<0.001; 3 months-6 months: 0.139 [-1.76,0.45], p= 0.8; HOA: Baseline-3 months 4.73 [4.133,5.332] p<0.001; 3 months-6 months: 0.112 [-0.512,0.736], p= 0.9).

Correlations between pain measure types pre-surgery were significantly positive in both OA groups, generally stronger in KOA groups than HOA groups, although these differences were relatively small. At 3- and 6-months post-surgery, the strength of the correlation of pain measures in the HOA group correlations were maintained; however, for the KOA, there was a strengthening of the correlations from baseline.

### 3.3 Pre-surgical pain levels do not relate to post-surgical pain relief

For all 4 pain outcome measures, we examined the relationship between pre-surgical pain and residual pain after surgery (100% residual pain meaning no change; 0% residual pain rendering complete relief), both for KOA and HOA at 3- and 6-months following surgery **(Figure 3)**. We observed no significant correlation between pre-surgical pain intensity and residual pain, indicating that pre-surgical pain levels are not useful predictors of relative change of pain with the procedure.

**Figure 3.**
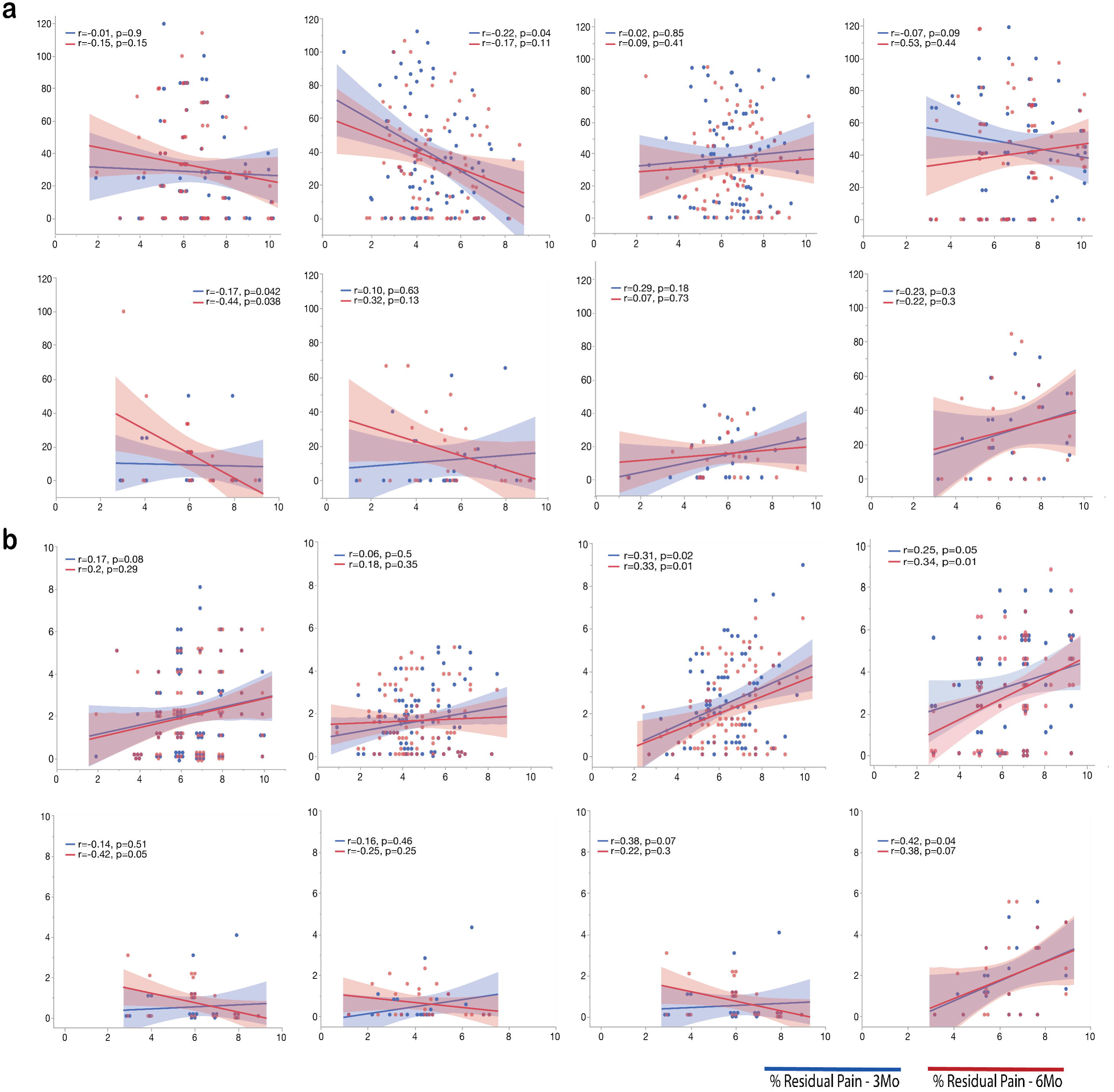
Influence of baseline pain intensity levels on post-surgical residual pain. The scatterplots depict patients’ percentage residual pain after surgery relative to pre-surgical levels (% residual pain, where 100% = no change from pre-surgical levels, 0% = full recovery), as a function of pre-surgical levels, for all four pain intensity measures. KOA (**a**) and HOA (**b**), at 3 (blue) and 6 (red) months post-surgery. Spearman correlations were not statistically significant, indicating that pre-surgical pain intensity did not influence post-surgery outcomes. Additionally, best fit slopes were not different between 3- and 6-months post-surgery, illustrating that surgery outcomes were essentially similar between 3 and 6 months. Symbols represent subjects. Shaded areas indicate 95% confidence intervals. BPI Severity, Brief Pain Inventory Pain: severity subscale; HOOS Pain, Hip Injury and Osteoarthritis Outcome Score: pain subscale; NRS, Numeric Rating Scale; SF36 Pain, Short-form (36) Health Survey: pain subscale.

### 3.4 OA related dimensions

Considering the broad battery of questionnaires and clinical measures collected, we sought to use a data dimensionality reduction approach to define dominant behavioral/clinical factors underlying OA pain. To this end, we applied a PCA analysis to the questionnaires and physical performance tests at baseline, focusing on the larger group of KOA patients (n=94). Pain intensity-related subscales were not included in this analysis, as they are the outcome measures to be modeled by PCA results. The correlations, organized by PCA results, are presented in **Figure 4a**. PCA identified 5 orthogonal components with eigenvalues>1.0, altogether explaining 69.9% of the variance. Given the observed loadings, we labeled them as: 1) *Affect*, composed of anxiety and depression subscales of HADS; *2) Pain Catastrophizing*, its highest factor loadings were the three maladaptive dimensions rumination, magnification and helplessness of PCS; 3) *Pain Quality*, dominated by the MPQ-sensory subscale and DN4, with high loadings regarding knee symptoms, knee related quality of life and sports and recreational ability; 4) *Health*, which was dominated by the SF-36 measures that quantify health status and health related quality of life; 5) *Physical Performance*, included high negative loading for 6MWT and high positive loading for TUG (**Figure 4b**). Note the five factors approximate the distinct domains surveyed by the questionnaires and tasks: HADS, KOOS, PCS, SF-36, and the combination of TUG and 6WMT. These five factors were used in subsequent model building to predict pain and residual pain.

**Figure 4.**
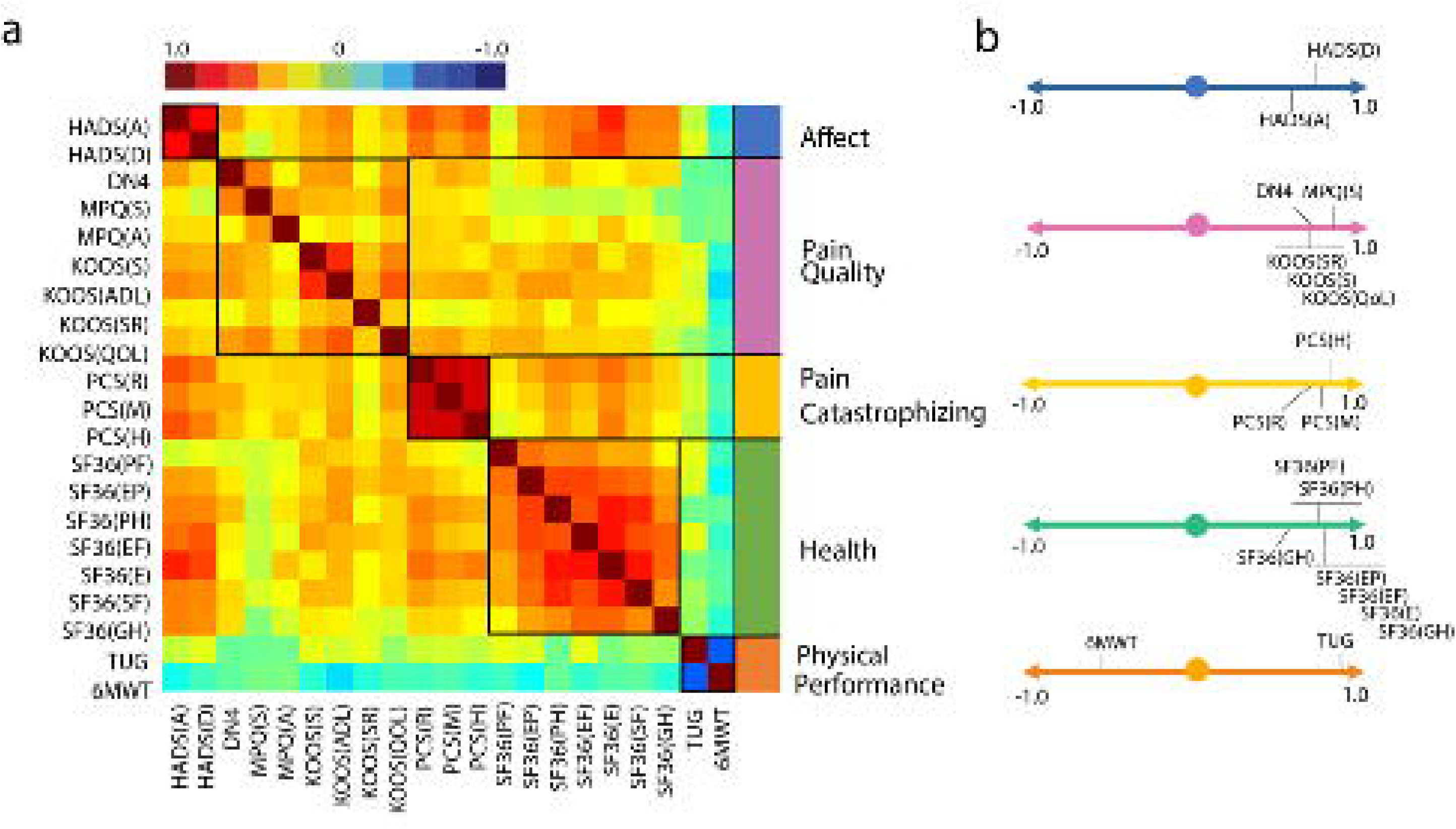
Principal component analysis identified five factors characterizing baseline KOA. Pain- and affect-related questionnaires, their subscales, and performance measures (prior to surgery) were examined together to identify dominant underlying factors. **a**. Correlation matrix ordered based on principal component analysis results (Pearson’s r represented by color bar). The five identified components were labeled according to membership properties. **b**. Factor loadings are shown for the five components. 6MWT, six minute walking test; DN4, The Neuropathic Pain 4 questions; HADS(A), The Hospital Anxiety and Depression Scale, Anxiety; HADS(D), The Hospital Anxiety and Depression Scale, Depression; KOOS, Knee Injury and Osteoarthritis Outcome Score, (ADL – Function in daily living), (S -Knee Symptoms), (SR – Function in sport and recreation), (QOL – knee related quality of life); MPQ, McGill Pain Questionnaire, (A – Affective score) (S – Sensory score); PCS, Pain Catastrophizing Scale, (R – Rumination subscale), (M – Magnification subscale), (H – Helplessness subscale); SF36, Short-form (36) Health Survey, (PF – Physical Functioning), (PH – physical role functioning), (EP – emotional role functioning), (EF – energy/fatigue), (E – emotional well-being), (SF – social functioning), (GH – general health); TUG, Timed -up and go test.

### 3.5 Modelling pain and TJR pain outcomes in OA

Next, we sought to model OA pain, using multi-factorial regressions (including only parameters that survived both forward and backward elimination), both at baseline and after surgery. Independent variables entered in our models are the five factors from the PCA results, together with relevant clinical/demographic variables: age, gender, educational level, body mass index, pain duration, and radiographic severity of OA.

#### 3.5.1 Pre-surgery KOA pain is defined by its quality, across pain measures

Pre-surgery KOA pain could be successfully modeled for all four outcome measures **(Table 3)**. All models reached statistical significance and accounted for 22-57% of variances of pain intensity. Pain quality emerged as the common dominant factor accounting for higher pain intensity throughout all scales. For NRS it was the only factor present in the model, whereas for the other 3 outcomes, additional factors were identified. BPI severity was predicted by higher levels of Pain Catastrophizing, KOOS Pain by worse Physical Performance and SF-36 pain by worse Health Status.

**Table 3.** Multiple regression models for KOA pain intensity at baseline for four different pain intensity measures.

| Model | b | SE | β | t | p | Adjusted R<sup>2</sup> |
| --- | --- | --- | --- | --- | --- | --- |
| BPI Pain Severity |  |  |  |  |  |  |
| Pain Quality | 2.526 | .670 | .385 | 3.770 | .000 | .275**<br>F(2,92)=18.816 |
| Pain Catastrophizing | 1.430 | .639 | .229 | 2.239 | .028 |  |
| NRS |  |  |  |  |  |  |
| Pain Quality | 0.851 | 0.162 | 0.479 | 5.258 | .000 | .229**<br>F(1,93)=27.652 |
| KOOS Pain |  |  |  |  |  |  |
| Pain Quality | 11.499 | 1.126 | .713 | 10.216 | .000 | .573**<br>F(1,92)=63.379 |
| Physical Performance | 2.426 | 1.123 | .151 | 2.160 | .033 |  |
| SF36 Pain |  |  |  |  |  |  |
| Health | 10.553 | 1.514 | .602 | 6.970 | .000 | .513**<br>F(2,92)=50.504 |
| Pain Quality | 3.438 | 1.573 | .189 | 2.186 | .031 |  |

KOA baseline pain intensity was explained by different pain-related characteristics, depending on the questionnaire used to capture pain intensity. While the pain quality factor was incorporated in all four regression models, additional unique influences were also identified for three of the four pain intensity scales. Displayed statistics are from the final step of each model. **b**, unstandardized regression coefficient; **SE**, standard error; **$\beta$** , standardized regression coefficient; **F**, obtained F-value; **t**, obtained t-value; **R<sup>2</sup>**, proportion of variance explained. \* $p \leq 0.05$ , \*\* $p \leq 0.01$ . Displayed statistics are from the final step for each dependent variable. BPI Severity, Brief Pain Inventory Pain: severity subscale; HOOS Pain, Hip Injury and Osteoarthritis Outcome Score: pain subscale; NRS, Numeric Rating Scale; SF36 Pain, Short-form (36) Health Survey: pain subscale.

#### 3.5.2 Models predicting pain intensity and residual pain after surgery in KOA

Next, we sought to model absolute pain intensity after surgery and residual pain (reflecting within subject change from pre-surgery) for all four pain measures, using the parameters collected prior to surgery, thus searching for pre-surgery influences on post-surgical pain. Modeling was restricted to pain at 6 months post-surgery, since there were minimal differences between post-surgery pain at 3 and 6 months.

Results, **(Table 4)** demonstrated that only three of the four outcome measures for absolute post-surgical pain could be modeled, accounting for 0−24% of the variance, and obtained models were distinct for each pain measure. We obtained similar results when modeling residual pain 6 months post-surgery. Only three of the four pain measures could be modeled, accounting for even smaller 0-11% of the variance, and obtained models were distinct for each outcome measure, as well as from the models obtained for post-surgical pain. Note that obtained results seem paradoxical. Correlations between the four pain outcome measures increases post-surgery yet obtained, pre-surgery based, models diverge from each other, both for pain and for residual pain.

**Table 4.** Multiple regression models for post-surgical KOA pain intensity, and for percentage residual pain at 6-months post-surgery, for four different pain intensity measures.

| Post-surgical Pain Intensity |  |  |  |  |  |  |
| --- | --- | --- | --- | --- | --- | --- |
| Model | b | SE | β | t | p | Adjusted R<sup>2</sup> |
| BPI Pain Severity |  |  |  |  |  |  |
| Affect | 2.845 | .670 | .408 | 4.246 | .000 | .238**<br>F(2,82)=13.952 |
| Pain Duration | .277 | .096 | .278 | 2.893 | .005 |  |
| NRS |  |  |  |  |  |  |
| No predictive model – no variables entered in the equation. |  |  |  |  |  |  |
| KOOS Pain |  |  |  |  |  |  |
| Affect | 5.688 | 2.229 | .284 | 2.522 | .013 | .234**<br>F(3,81)=9.338 |
| Gender | 9.756 | 4.320 | .221 | 2.259 | .027 |  |
| Health | 4.246 | 2.026 | .231 | 2.060 | .043 |  |

| SF36 Pain |  |  |  |  |  |  |
| --- | --- | --- | --- | --- | --- | --- |
| Health | 7.032 | 2.292 | .310 | 3.068 | .003 |  |
| Gender | 15.176 | 5.520 | .278 | 2.749 | .007 |  |
|  |  |  |  |  |  | .196* |
|  |  |  |  |  |  | F(2,82)=9.730 |
| % Residual Pain |  |  |  |  |  |  |
| Model | b | SE | $\beta$ | t | p | Adjusted R<sup>2</sup> |
| BPI Pain Severity |  |  |  |  |  |  |
| Pain Duration | 1.413 | .536 | .274 | 2.635 | .01 |  |
| Health | 7.681 | 3.451 | .232 | 2.226 | 0.029 |  |
|  |  |  |  |  |  | .114** |
|  |  |  |  |  |  | F(2,82)=6.267 |
| NRS |  |  |  |  |  |  |
| No predictive model – no variables entered in the equation. |  |  |  |  |  |  |
| KOOS Pain |  |  |  |  |  |  |
| Physical Performance | 9.276 | 3.036 | .321 | 3.055 | .003 |  |
|  |  |  |  |  |  | .092** |
|  |  |  |  |  |  | F(1,83)=9.335 |
| SF36 Pain |  |  |  |  |  |  |
| Gender | 24.210 | 8.088 | .316 | 2.993 | .004 |  |
|  |  |  |  |  |  | .088** |
|  |  |  |  |  |  | F(1,83)=8.959 |

Different explanatory models were elicited for absolute versus relative (% residual pain) pain intensity after surgery, with the variance explained by each model being overall lower for % residual pain than that for absolute pain intensity. Again, explanatory variables were also distinct considering the four different outcome measures.
**b**, unstandardized regression coefficient; **SE**, standard error; **$\beta$** , standardized regression coefficient; **F**, obtain F-value; **t**, obtained t-value; **R<sup>2</sup>**, proportion variance explained. Gender: male coded as 0, female coded as 1. \* $p \leq 0.05$ , \*\* $p \leq 0.01$ . Displayed statistics are from the final step for each dependent variable.
BPI Severity, Brief Pain Inventory Pain: severity subscale; HOOS Pain, Hip Injury and Osteoarthritis Outcome Score: pain subscale; NRS, Numeric Rating Scale; SF36 Pain, Short-form (36) Health Survey: pain subscale.

#### 3.5.3 Do KOA models of pain and residual pain generalize to HOA?

Given the smaller data available in HOA (n=22), and the large number of independent variables and four pain outcome measures, we limited HOA modeling. We only tested the extent to which models obtained in KOA are shared with HOA. Therefore, regression models were constructed for HOA pre-surgical pain, 6-months absolute post-surgical pain and residual pain using only parameters identified for KOA. Pre-surgery, the multiple regression successfully modeled pain intensity for HOOS Pain (equivalent to KOOS pain), F (2,22) =24.308, p<0.005, however only one of the two variables entered, Pain Quality, was significant (β=.764, p=0.005). For SF-36 pain, the model obtained for KOA was also applicable, F (2,22) = 23.55, p<0.001. Here the factor Health (β=.732, p<0.001), but not Pain Quality was significant. NRS and BPI in HOA failed to be modeled by the HOA parameters.

Pain and residual pain for HOA after surgery, failed to be modeled by KOA parameters, for any of the four pain scales.

### 3.6 A composite measure of pain intensity

As the pre-surgical parameters predicting post-surgical pain or residual pain for four pain outcome measures captured distinct independent variables, we reasoned that each may be reflecting specific characteristics and thus combining all four measures would predict larger variance and incorporate the component characteristics. Therefore, we constructed the composite, average score, of all four pain outcome measures and studied its properties. Similar to the trend set by the unitary scales, aggregated pain intensity levels were similar between OA groups pre-surgery (mean of ∼6, 0-10 scale), decreased after surgery (∼2-1), to a greater extent for hip than for knee, and was stable between 3 and 6 months **(Figure 5a)**. The distribution of residual pain with the aggregate measure again highlighted better surgical outcomes in HOA: 74% of KOA patients reported at least 20% reduction of the initial pain, but only 46% of HOA. The number of patients sustaining higher pain levels decreases for both OA groups, more dramatically for hip than for knee OA **(Figure 5b)**.

**Figure 5.**
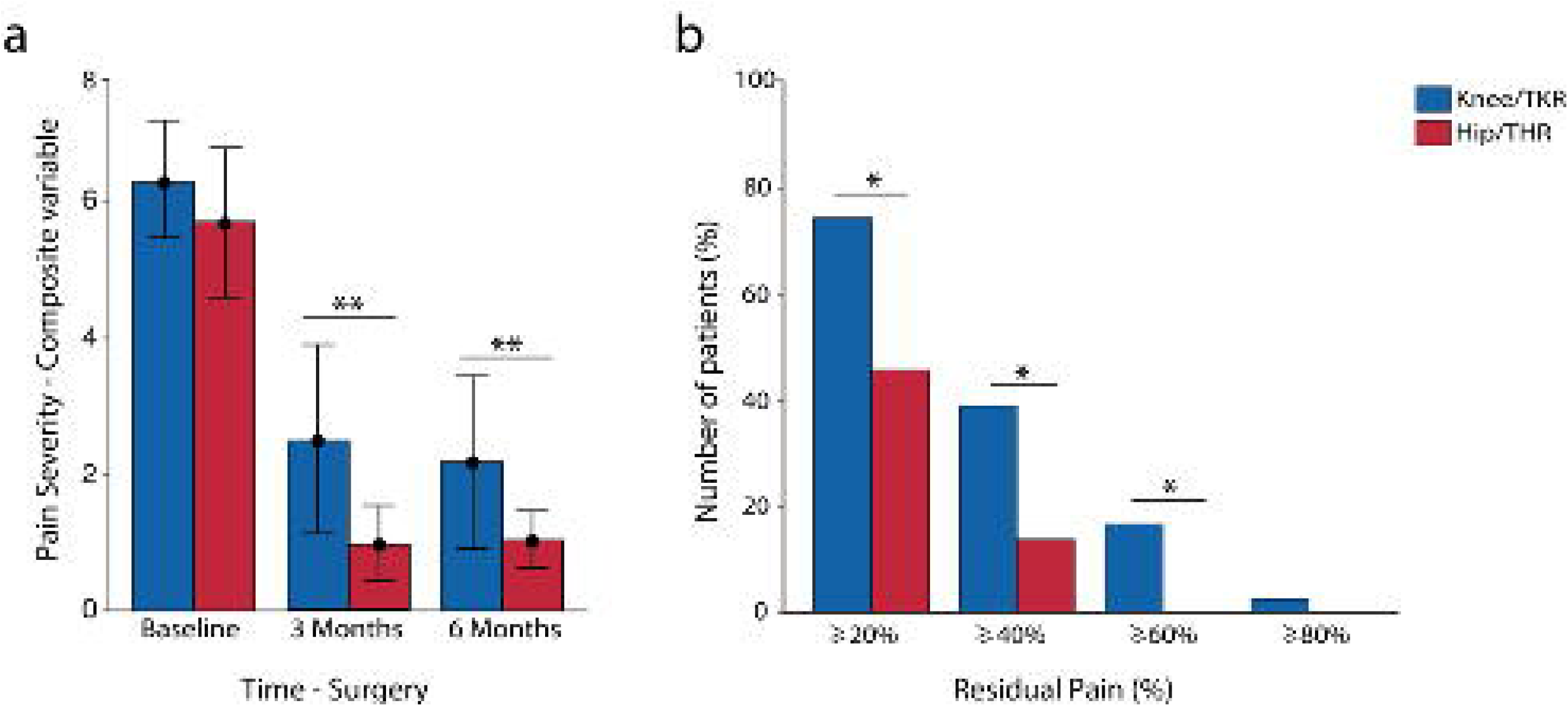
Composite pain intensity variable defined severity, and residual pain after surgery. Aggregated variable of pain intensity created by averaging the four pain outcome measures. a) - Pain intensity for KOA (blue) and HOA (red) at baseline, and at 3 and 6 months after surgery. There was an interaction between condition (KOA and HOA) and time on the levels of pain intensity (two-way mixed ANOVA, F (2,208) =3.67, p=0.03). Pain intensity after surgery was higher for KOA, both at 3 months (F (104) =17.65, p<0.001) and 6 months (F (104) =10.037, p<0.001). For each bar, circles represent the mean value, and bars indicate standard deviation. b) The bar graph depicts the percentage of patients (%) at each category of extent of residual pain at 6 months post-surgery. KOA patients sustain higher levels of pain relative to HOA (20%: X^2^ (1) =6.43, p=0.011; 40%: X^2^(1) =4.71, p=0.03; 60%: X^2^(1)=3.88, p=0.049). **p<0.01; *p<0.05.

Next we tested how pre-surgical factors predict 6-months post-surgical KOA pain, using our aggregate measure **(Table 5)**, again modeling pain and residual pain for KOA patients. For post-surgical aggregated pain severity, the model explained 19% of the variance and included worse health state, lower degree of structural articular damage, and poor results in the physical performance tests. For residual pain, the model explained 7% of the variance and Physical Performance was the only predictive factor. Using these variables to predict HOA post-surgical pain and residual pain we could not find any statistically significant models.

**Table 5.**
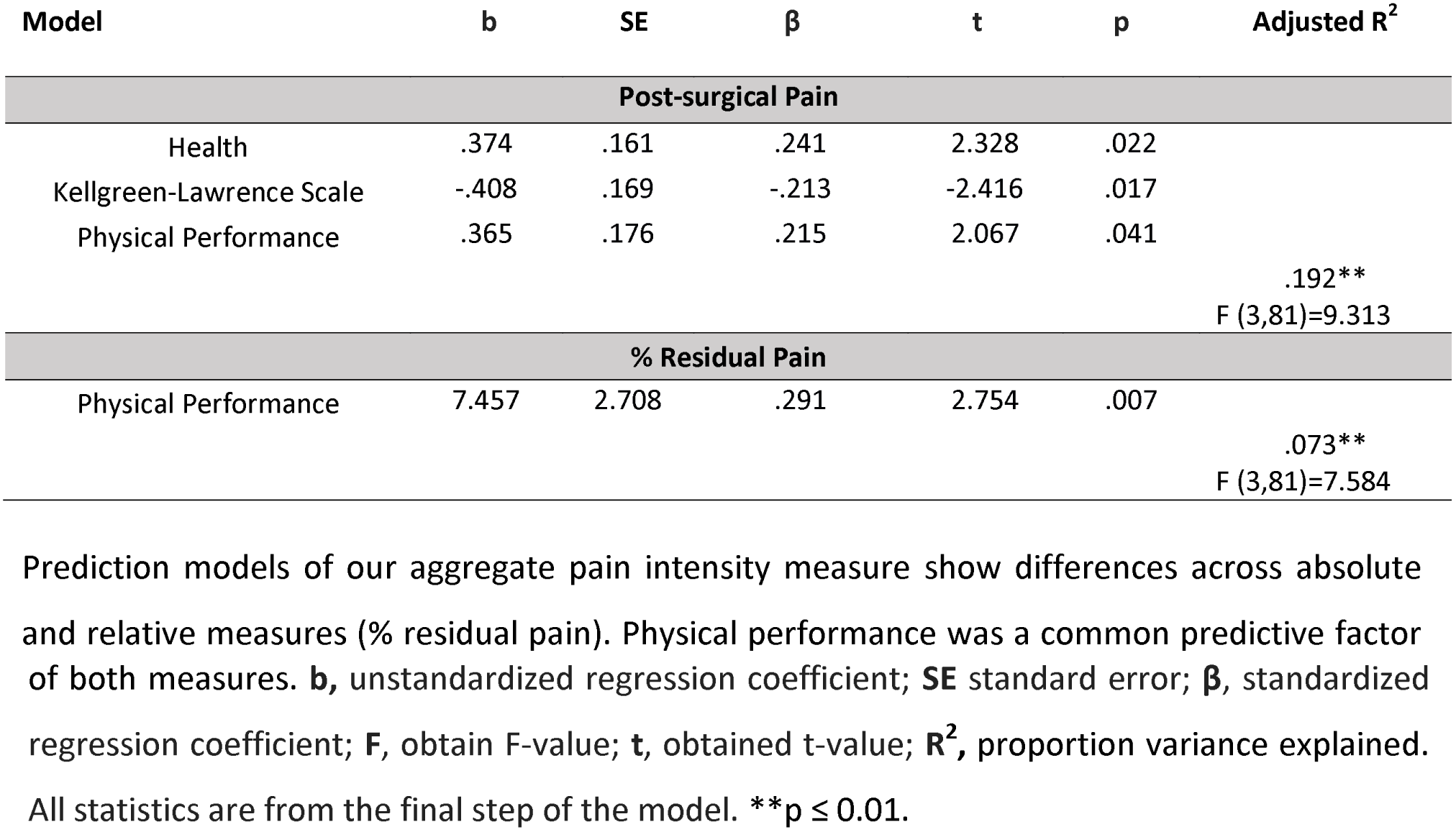
Multiple regression analysis for KOA pain intensity and % residual pain at 6-months post-surgery, using our aggregated variable for pain intensity (average of four pain intensity questionnaire measures).

| Model | b | SE | $\beta$ | t | p | Adjusted R<sup>2</sup> |
| --- | --- | --- | --- | --- | --- | --- |
| Post-surgical Pain |  |  |  |  |  |  |
| Health | .374 | .161 | .241 | 2.328 | .022 | .192**<br>F (3,81)=9.313 |
| Kellgreen-Lawrence Scale | -.408 | .169 | -.213 | -2.416 | .017 |  |
| Physical Performance | .365 | .176 | .215 | 2.067 | .041 |  |
| % Residual Pain |  |  |  |  |  |  |
| Physical Performance | 7.457 | 2.708 | .291 | 2.754 | .007 | .073**<br>F (3,81)=7.584 |

Prediction models of our aggregate pain intensity measure show differences across absolute and relative measures (% residual pain). Physical performance was a common predictive factor
of both measures. **b**, unstandardized regression coefficient; **SE** standard error; **β**, standardized regression coefficient; **F**, obtain F-value; **t**, obtained t-value; **R<sup>2</sup>**, proportion variance explained. All statistics are from the final step of the model. **\*\*p** ≤ 0.01.

### 3.7 Network analysis of pain dimensions

An alternative to regression-based modeling of the effects of TJR on OA pain is to examine properties of the correlation matrix identified pre-surgery (**Figure 4**) as a function of type of OA and time from surgery. Representing such correlation matrices as networks provides insights regarding organizational topography and changes in the inter-relationships between pain characteristics that define the OA state. Therefore, we calculated these networks pre-surgery, and three- and six-months post-surgery (**Figure 6**).

**Figure 6.**
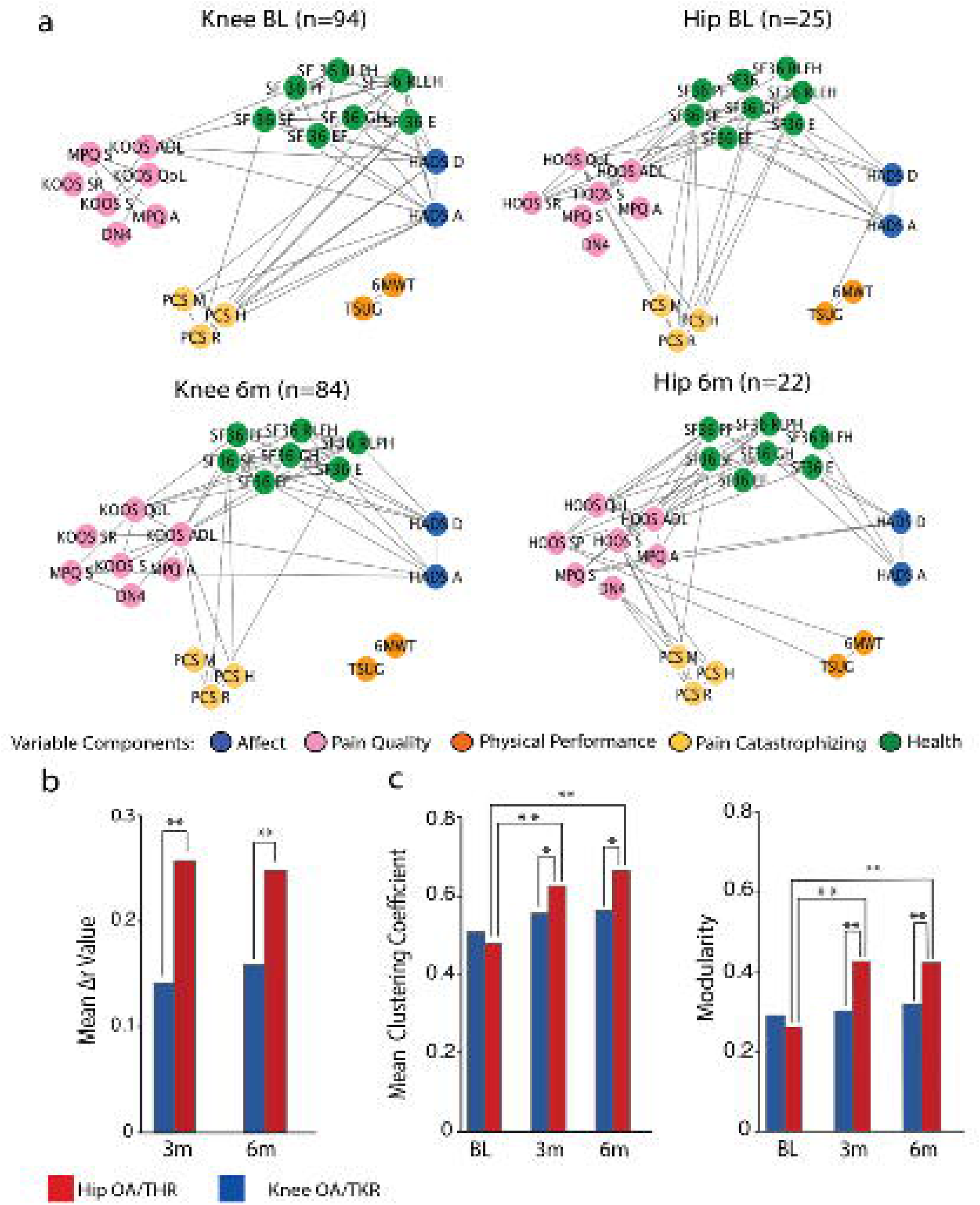
Network representation of OA pain characteristics. **a)** Network graphs depict interrelations between clinical and pain-related questionnaire subscale measures at baseline, and at 6 months post-surgery, for KOA and HOA patients. Network communities were derived from the PCA analysis. Links represent the top 25% correlations of each network. **b)** The bar graph displays mean change of global correlation coefficients (Pearson’s Δr) for KOA and HOA, at 3- and 6-months post-surgery. Both groups had significant change in the overall interrelations between clinical and pain-related characteristics (KOA mean Δr 3months: 0.14, t=13.37, mean Δr 6months: 0.16 t=14.93, HOA mean Δr 3months:0.28, t=8.72, mean Δr 6months:0.26 t=9.23, p<0.001). The extent of change remained stable from 3 to 6 months post-surgery and was substantially higher in the HOA group at 3 months (t=4.62, p<0.001) and 6 months (t=3.44, p<0.001). c) Graph theory-based modularity and mean clustering coefficients for correlation networks at baseline, 3 and 6 months. The HOA networks shows significant topological reorganization 3 months (mcc: t=-8.19, modularity: t=-9.22, p<0.001) and 6 months after surgery (mcc, t=-10.62, modularity, t=-9.02, p<0.001), while KOA remains stable. BL, baseline; 3m, 3 months; 6m, 6 months; Statistical risk probability was computed under 10.000 times repeated random resampling. **p<0.001, *p<0.05.

Regarding the pre-surgery KOA network, factors Affect, Pain Catastrophizing and Health presented salient edges (significantly high correlations) among them. Pain Quality showed a lower number of edges connecting with other factors (only through subscale KOOS-ADL). Physical performance was segregated from the other factors. For HOA, Affect and Pain Catastrophizing did not share any salient correlations. Pain Quality was highly correlated to Health and to a lesser extent to Pain Catastrophism. Physical performance was again segregated.

At six months after surgery topological differences were identified in both KOA and HOA groups. For the KOA network, Affect and Pain Catastrophizing no longer presented salient edges. Pain quality shared a higher number of edges with Affect and Health. Physical Performance continued to be isolated, sharing no edges with other components. For HOA, Pain Catastrophizing lost its prominent edges with Health and was only linked with Pain Quality. Physical Performance showed links with one variable in Pain Quality (HOOS Sports and Recreational) (**Figure 6a**).

To quantify topological changes in these network architectures we derived network measures and compared them between groups and as a function of time. We calculated change in strength of connectivity (change in correlation coefficients for all pairs of subscales, Δr-value) both for 3- and 6-months post-surgery. For further comparison intra- and inter-groups, we computed statistical probability using 10,000 permutations with random resampling.

Inside each group, there was a significant change in Δr-value, for both KOA and HOA, at 3 and 6 months, with no differences between 3 and 6 months in each group, indicating that post-surgical connectivity is stable in time. When comparing between KOA and HOA groups, connectivity change was larger for HOA both at 3 and 6 months (**Figure 6b**).

Lastly, we evaluated the clustering coefficient and modularity of the networks and assessed differences between groups. For both measures, KOA networks remained stable after treatment. HOA, on the other side, showed a significant change in both measures, from baseline to 3 and 6 months. From 3 to 6 months the networks remained stable (**Figure 6c**).

Overall, we observed that pain characterizing networks for KOA and HOA are quite distinct from each other prior to surgery while displaying similar topology, and only the HOA network is significantly reorganized post-surgery.

## Discussion

This study examined KOA and HOA pain prior and after TJR surgery. We used a systematic and structured approach, together with data reduction techniques, to investigate the properties of OA pain, its change with surgery, and factors that influence these measures. By using four distinct pain intensity quantifying measures, two distinct types of joint OA, and measures collected at pre-, 3, and 6 months post-surgery, we examine the contribution of a large number of potential influences, many of which have been reported to be risk factors for OA pain persistence post-TJR. As available data were larger for KOA, we performed model building in this group and tested identified variables in HOA. Each of the four-pain intensity measures we used, either alone or their aggregate, demonstrated an overall decrease in OA pain after surgery in both OA groups. For HOA, these decreases were at least twice as large and in a larger proportion of patients, as residual pain >60% was observed only in 18% of KOA and 0% of HOA. A striking and perhaps unexpected result was how little OA pain changed from 3- to 6-months post-surgery in both groups. Neither the mean pain nor pain characteristics, as assessed by network properties, showed any important changes over this time period, although large changes were seen between pre-surgery and 3-months post-surgery. Our regression models showed that commonly assessed clinical and behavioral measures prior to surgery fail to reliably predict pain outcomes in knee and hip OA patients.

OA pain and persistent pain after TJR have been previously studied using multiple pain outcome scales. These can be divided in two major groups, general measures such as NRS, visual analog scale [32], SF-36 bodily pain and BPI pain severity, and OA specific measures as Western Ontario and McMaster Universities Osteoarthritis Index (WOMAC), KOOS/HOOS pain score and the Oxford Knee Score pain subscale [7, 33]. Such studies suggest that different pain outcomes relate to different facets of the pain experience in knee OA [34]. Using four different pain intensity outcomes, three of them from the category of general pain scales and one specific for OA (HOOS/KOOS), our results show that although correlations between these measures are positive and mostly significant (both at baseline and post-surgery), BPI pain severity tends to underestimate pain intensity and SF-36 pain tends to overrate pain intensity after surgery both in KOA and HOA groups. Still, all four measures decreased 3-months post-surgery, and all remained unchanged between 3- and 6-months post-surgery.

Pain outcomes concerning persistency are commonly studied using the absolute value of intensity after surgery, or dichotomizing the outcome using a fixed threshold that varies across studies [35-38]. This implies that the treatment has a constant effect. A change may show the health improvement in a more observable way. Here we chose to use percent change. It evaluates the within subject effect of treatment and is a better metric for assessing the influence of TKR on post-surgical OA pain. All individual residual pain score showed independence from baseline pain, implying pain relief post-surgery does not depend on entry scores.

An important remark concerning post-surgical pain and its risk prediction is that it relies on how it is defined and thus also on how one measures the pain outcomes. Chronic post-surgical pain is accepted as the pain that persists at least three months after surgery, different in characteristics from pre-operative pain, and without other causes such as infection or technical failure [5]. Our results are generally consistent with this definition and further advance the concept. Firstly, we observe that models characterizing OA pain at baseline do not generalize to pain post-surgery. Second, the amount of variance explained with the regression models for pain intensity decreased from pre-surgery (accounting for 23-57% of pain intensity variance), to post-surgery (accounting for 20−24% of variance), and further decreased when modeling residual pain (accounting for 9-11% of variance). Given that residual pain is a more direct measure of the influence of the surgical intervention than the absolute value of pain intensity, our models at best could only explain 11% of the variance of the surgery related OA pain. Importantly, the models obtained for residual pain for each pain measure could not be replicated when modeling the aggregated measure, again attesting that these models are weak and inconsistent. Third, studies report that pre-operative OA pain intensity has a strong influence on post-surgical outcomes [7]. It was recently argued that the evidence for this influence is of low-quality [9], and our results support the failure of pre-operative pain as a predictor of post-surgical outcomes. If we consider residual pain as the more direct metric of the influence of surgery on post-surgical OA pain, then we observe no relationship between pre-surgery OA pain and residual pain. Fourth, the network analysis shows large changes in the interrelationships between pain related characteristics post-surgery. Thus, our analysis, especially for KOA where we examined multiple models, suggests that the post-operative pain is minimally related to the pre-operative pain properties. Our results in HOA, although not as strong, are also consistent with this notion.

Given the small sample size in HOA, we limited the models and statistical tests in this group. Models derived from KOA did not generalize to HOA. Thus, HOA pain models remain to be studied in larger data sets in the future, and with additional parameters not included here. However, our results repeatedly confirm that pain relief is better in this group and this is accompanied with larger changes in the network properties. We observed larger changes in clustering coefficient and in modularity in HOA, implying that the pain personality in HOA is being fractured with pain relief, rendering different factors independent from each other. These findings are all consistent with earlier reports showing that the improvement in pain and physical function after arthroplasty is greater for hip than knee OA [3], even though symptomatic presentation of HOA is associated with more advanced radiological disease [39]. Determinants for persistent pain after THR are less studied, and evidence is limited and conflicting [40]. The full scope of the differences in TJR outcomes between both conditions requires further studies.

A major focus of this paper was to find predictors for pain and pain persistency, and although we show that there is a high variability concerning scales and outcome definitions, some of the findings deserve further discussion. At baseline, we observed that across the four scales and the aggregated pain measure, Pain Quality (constituted mainly by neuropathic pain profile and sensory quality of MPQ) related to higher pain intensity. When we modelled residual pain after surgery, each scale unveiled different predictors. No homogeneous result could be retrieved. For our aggregated variable, residual pain could be predicted by a different parameter, while the expectation was that this aggregated measure would capture larger variance for the same factors. A previous study has reported that worse pre-operative functional status was related to pain persistency [41] the present results do not confirm the role of pre-operative walking ability on post-operative KOA pain.

It has been reported that the greatest improvement in patients undergoing TJR happens in the first 3 months after surgery [3]. Although a precise timeline for pain recovery is difficult to draw, our results support the finding that pain persistence at 3 months should be regarded as a serious caveat for longer-term post-surgical pain.

An important weakness of the present study was the imbalance of available data between KOA and HOA. Moreover, there were important demographic differences between the two groups which could not be corrected for due to the limited available sample in HOA. Thus, we cannot rule out the influence of these factors on the models derived from KOA and tested in HOA. The total number of subjects included in this study is small, however not a limitation for the statistical modelling applied. Regarding our sample characteristics, patients were enrolled in the same center, and the population included is ethnically homogeneous, thus caution is needed in generalizing the present’s study results to other populations. The follow-up time was limited to 6 months, what can also be regarded as a limitation. Research for longer term follow-ups, with larger numbers of subjects conveying a similar multimodal approach should be tested in the future. We also did not collect multiple measures that in the literature have been suggested to influence both baseline pain and post-surgical pain. For instance, measures of widespread hypersensitivity, temporal summation of pain and impaired endogenous pain inhibition assessed by quantitative sensory testing, have been suggested to contribute to poor pain relief following TKR [42, 43], however see [44]. It was also previously shown that OA patients present central nervous system structural and functional maladaptive changes [45, 46]. We will test the latter concept in this same group of participants using their brain imaging results.

In conclusion, our results show distinct pain scales relate to different aspects of the pain experience. Post-surgery residual pain scores show independence from baseline pain, there is a reorganization of pain related biopsychosocial parameters with surgery, however, an unpredictability of post-surgical OA pain from pre-surgical pain related parameters.

## Acknowledgments

We thank to Dr. Paulo Oliveira, for the clinical support and supervision of patients enrolled in the study and all the members of the Orthopedic Department of Centro Hospitalar Universitário de São João, Porto, for their support with data collection. Dr. Lina Melão and Dr. Patricia Leitão for the assistance in the radiographic classification. We are thankful to all Apkarian lab and Galhardo lab members that contributed to this study with their time and resources.

## Disclosures

J.B. was funded through CCDRN [Norte-08-5369-FSE-000026], OARSI Collaborative Scholarship 2018 and Luso-American Development Foundation R&D@PhD scholarship grant. This research did not receive other specific funding from agencies in the public or commercial sectors.

## Notes

#### Summary of Updates

Figure order

